# PIEZO ion channel is required for root mechanotransduction in *Arabidopsis thaliana*

**DOI:** 10.1101/2020.08.27.270355

**Authors:** Seyed A. R. Mousavi, Adrienne E Dubin, Wei-Zheng Zeng, Adam M. Coombs, Khai Do, Darian A. Ghadiri, Chennan Ge, Yunde Zhao, Ardem Patapoutian

## Abstract

Plant roots adapt to the mechanical constraints of the soil to grow and absorb water and nutrients. As in animal species, mechanosensitive ion channels in plants are proposed to transduce external mechanical forces into biological signals. However, the identity of these plant root ion channels remains unknown. Here, we show that *Arabidopsis thaliana* PIEZO (AtPIEZO) has preserved the function of its animal relatives and acts as an ion channel. We present evidence that plant PIEZO is highly expressed in the columella and lateral root cap cells of the root tip which experience robust mechanical strain during root growth. Deleting PIEZO from the whole plant significantly reduced the ability of its roots to penetrate denser barriers compared to wild type plants. *piezo* mutant root tips exhibited diminished calcium transients in response to mechanical stimulation, supporting a role of AtPIEZO in root mechanotransduction. Finally, a chimeric PIEZO channel that includes the C-terminal half of AtPIEZO containing the putative pore region was functional and mechanosensitive when expressed in naive mammalian cells. Collectively, our data suggest that *Arabidopsis* PIEZO plays an important role in root mechanotransduction and establishes PIEZOs as physiologically relevant mechanosensitive ion channels across animal and plant kingdoms.

## Main

Plants extend roots within the soil to access water and nutrients as well as provide stability for the aerial parts of the plant. Underground barriers caused by drought and/or heterogeneous soil components can exert mechanical resistance that alters root extension and penetration^1–3^. The root cap at the very tip of the primary root is a dynamic organ that includes different classes of stem cells which divide asymmetrically and is essential for growth through harder media and soils^4^. Bending or poking root tips elicits a transient Ca^2+^ influx with short latency that is blocked by lanthanides including Gd^3+^, a non-selective inhibitor of mechanically-activated (MA) cation channels^5–7^. However, the molecular identity of putative ion channels underlying this response is unknown. Only a few mechanosensitive ion channels have been described in plants^8^. MSL8, plays a mechanosensory role in pollen^9^, MSL10 is involved in cell swelling^8,10^, and OSCA1 has mainly been characterized for its role in osmosensation^11^. It has been proposed that MCA1, expressed in the elongation zone but not the root cap, is a stretch-activated calcium permeable ion channel involved in soil penetration; however, evidence for its being a bona-fide ion channel capable of detecting mechanical force is lacking^12–14^. The genome of *Arabidopsis thaliana* encodes an ortholog of the mammalian mechanosensitive ion channels *PIEZO1* and *PIEZO2*^15^. Given that PIEZOs play prominent roles in multiple aspects of animal mechanosensation and physiology^16–19^, we investigated the role of AtPIEZO in plant mechanosensation. A recent study reported that AtPIEZO regulated virus translocation within the plant, but its specific role in mechanotransduction was not addressed^20^. Here we use genetic tools, electrophysiological methods and calcium imaging to investigate the role of AtPIEZO in root mechanosensation.

To localize the expression of AtPIEZO in *Arabidopsis*, we used *AtPIEZO* promoters fused to the reporter gene β-glucuronidase (GUS) and generated two *AtPIEZOpro::GUSPlus* constructs with different promoter lengths, 823 bp and 2000 bp. Both constructs showed similar GUS expression, with high levels observed in upper root, both primary and lateral root caps, and pollen grains (Fig.1a, c, and Extended Data Fig. 1). We also detected GUS activity in the root vasculature and in trichromes (plant hairs) (Fig.1b and Extended Data Fig. 1). Cross-sections of root tips revealed expression in lateral root cap (LRC) cells and columella cells (Fig.1g,h), that are thought to be important in detecting mechanical forces during root penetration into the soil^4^. When plants were grown inside Murashige and Skoog medium (0.5X MS; 0.85% agar (8.5 g/l)) rather than on top of it, higher GUS signal intensity was observed in the upper root and root cap of the seedlings suggesting expression is enhanced when mechanical stress is applied to roots (Fig.1d,e). This increase in GUS activity was confirmed by quantitative real time PCR; *AtPIEZO* expression was 3-fold higher in plants grown inside MS media (Fig. 1f).

**Figure 1.**
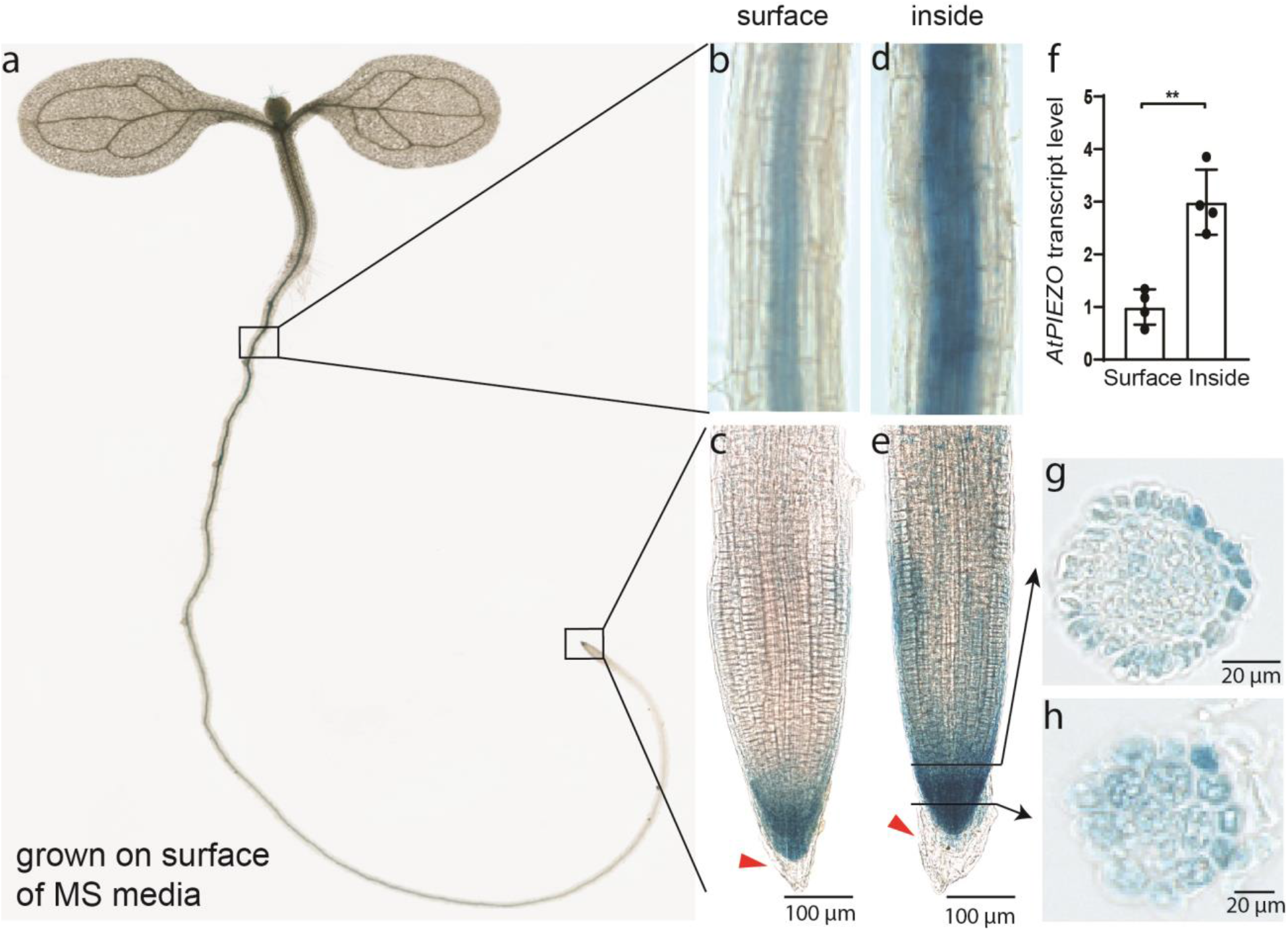
Expression pattern of *AtPIEZO::GUSPlus* reporter line in Arabidopsis root. **a**, Expression pattern of the GUS reporter protein under 2000 bp of the *AtPIEZO* promoter in a 7 days old seedling. **b**-**c**, Expression in the upper root and root tip when the plant is grown on the surface of standard MS media. Red arrowhead indicates that *AtPIEZO* is no longer expressed in the oldest root cap cells that are most distal and are known to be sloughed off. **d-e**, Expression in the upper root and root tip grown inside the MS media. Black arrows indicate the cross section of root displayed in panels g and h. **f**, qRT-PCR for *AtPIEZO* in upper root of plants grown on the surface (“surface”) or within the MS media (“inside”). \*\**P* < 0.01, N=4 (mean±s.d.). **g-h**, Cross sections of the root cap in GUS reporter lines which indicate expression in columella and LRC cells.

Next, to investigate the role of AtPIEZO in plant physiology and development, we generated two *piezo* CRISPR/Cas9 knockout mutant lines: one in which the entire gene was deleted (referred to as *piezo*-*FL*), the other in which the C-terminal half of the gene that encodes the putative channel pore based on its homology to mouse PIEZO1 (mPIEZO1) was deleted (referred to as *piezo*-*CT*). AtPIEZO has 27% amino acid identity with mPIEZO1 with similar overall topology and 38 predicted transmembrane domains (Extended Data Fig.2). We confirmed the lack of *AtPIEZO* transcripts in both mutants by PCR and RT-qPCR in samples harvested from the leaves as well as the roots (Extended Data Fig. 3). We did not observe any significant growth difference between WT and *piezo* mutants in roots or aerial parts when grown in MS media.

**Figure 2.**
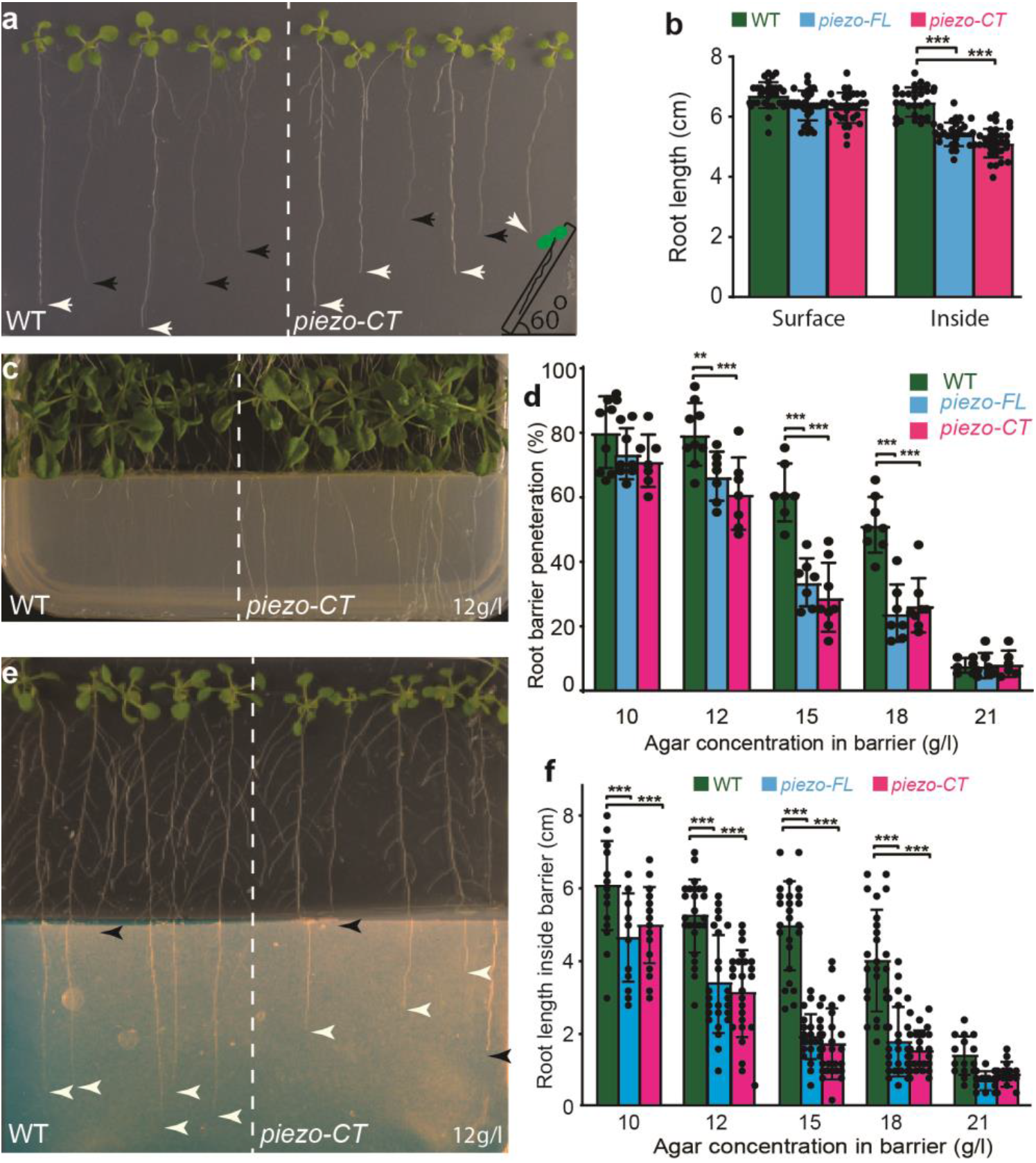
Penetration of roots into hard media is compromised in *piezo* mutants. **a**, Root length of *Arabidopsis* roots grown 9 days vertically at 60° to stimulate growth into the MS media. Black arrowheads indicate roots within the MS media, white arrowheads indicate root growth on the surface of MS media. **b**, Root length of WT and *piezo* mutants 9 days after germination (N=30-36; mean±s.d). **c**. Representative plate of 18 day old *Arabidopsis* seedlings challenged by root barriers. The more visible roots have not penetrated the barrier and are on the surface of the MS media. WT and mutants were grown vertically at 90° on normal MS media (0.5X MS + 8.5 g/l agar) before reaching the barrier (0.5X MS + 12 g/l agar). **d**, The percentage of roots that penetrate barriers of different concentrations of agar as indicated (n=8-11 plates) and each plate consists of 7-12 seedlings. **e**. Representative plate of 2 week-old *Arabidopsis* seedlings challenged by root barriers indicating the difference root growth length in the denser media (imaged brightness was adjusted for clarity). White arrowheads indicate roots within the MS media, black arrowheads indicate root growth on the surface of MS media or roots that did not penetrate the barrier. **f**, Root lengths of 2 weeks old seedlings that have penetrated the barriers (indicated) (N=25-28, mean±s.d.). \*\**P* < 0.01, \*\*\**P* < 0.001.

Based on the robust root expression of *AtPIEZO*, we sought to evaluate its role in root growth. We grew *A. thaliana* seeds on the surface of the MS media and plates were positioned vertically (at a 90° angle). The length of the seedling roots of WT and the two mutants were not different when grown on top of the MS media (Extended Data Fig. 4 a, b). However, when seedling roots grew within the MS media, mutant roots were shorter compared to WT. To confirm the differences observed for root penetration and growth inside the media, we challenged the roots at a 60° plate angle to stimulate growth into the media. Again, we observed that the roots of both mutants were shorter than WT roots (Fig. 2a-b). To further investigate root penetration, plants were grown at 60° in MS media containing different agar concentrations (7, 8.5 (standard), and 10 g/l) mimicking different levels of soil hardness. The root lengths of *piezo* mutants and WT plants were similar in the lowest agar concentration (7 g/l; Extended Data Fig. 4c). However, at higher agar concentrations (8.5 and 10 g/l), the average root lengths of both mutants were significantly shorter by about 17% and 18%, respectively, than observed for WT. These data show that mutants had shorter length than wild type in hard medium (Extended Data Fig. 4c).

To assess the response of roots to stiff materials that might be encountered during growth, we challenged roots with barriers of varying stiffness consisting of 10, 12, 15, 18 and 21 g/l agar in MS media (Fig. 2d). We plated *A. thaliana* seeds on the standard agar concentration in MS media (8.5 g/l agar), 2 cm above the barrier. Within 4-5 days after germination, seedling roots of all genotypes reached the barrier. At this point, three different scenarios were observed: 1) penetration across the barrier, 2) root coiling and delayed penetration after growing at the interface surface, or 3) no penetration (Extended Data Fig. 5c-d). At a 10 g/l barrier, 80% of WT roots penetrated the harder agar while only 74% and 73% of *piezo-FL* and *piezo-CT*, respectively, were able to penetrate (n=9). As the agar concentration increased, the barrier penetration phenotype in the mutants became more pronounced. For example, at 15 g/l, the penetration percentage for WT was 58% while only 29% of *piezo-FL* and 26% of *piezo-CT* roots penetrated (n=11) (Fig. 2d; Supplementary Video 1). Furthermore, mutants showed a delayed penetration with excessive coiling on the barrier surface (Extended Data Fig. 5d).

For the roots that penetrated the various barriers, root length inside the barriers was more variable and shorter in the mutants compared to WT (Fig. 2e,f). For example, at a 12 g/l agar barrier concentration, the root length of WT was 5.2±01.2 cm (n=28), while it was 3.4 ±1.3 cm and 3.1±1.2 cm for *piezo-FL* and *piezo-CT*, respectively (n=25-28) (Fig. 2f). The shorter root length in mutants became more severe in 15 or 18 g/l agar. Although mutant root coiling at the barrier interface delays root penetration and contributes to the decreased total root length, shorter roots are observed for mutants seeded directly into media containing 8.5 and 10 g/l agar (Extended Data Fig. 4c), indicating that the velocity of root growth is slowed in denser media.

These results implicate a role of AtPIEZO in mechanosensory processes in plants. To assess whether AtPIEZO is a mechanosensitive ion channel like its animal homologs, we cloned the full length coding sequence (7455bp) into a mammalian expression vector (see Methods). Transient heterologous expression of PIEZO proteins from various animal species (including mammals and flies) confer robust MA currents^15–17^. We heterologously expressed either native or codon-optimized *AtPIEZO* in HEK293T Piezo1 knockout (HEK P1KO) cells^21^. Neither native nor codon-optimized *AtPIEZO* revealed MA currents in two separate assays for mechanotransduction: poking cells with a fire-polished glass pipette^22^ and stretching the membrane at the tip of the pipette in cell-attached patch clamp recordings^22^. To determine whether the lack of response was due to improper trafficking to the plasma membrane, MYC-tags were inserted at five separate predicted extracellular loops of the protein based on homology between transmembrane domains of AtPIEZO and mPIEZO1^22–24^ (Extended Data Fig 2). Immunostaining with an anti-MYC antibody detected AtPIEZO expression in HEK P1KO cells, however, non-permeabilized staining revealed that AtPIEZO did not traffic to the membrane (Extended Data Fig. 6). We next generated chimeras between mPiezo1 and codon optimized AtPIEZO in an effort to traffic chimeras containing the putative pore domain of AtPIEZO to the membrane. As the pore region of PIEZOs is located at the C-terminus^22–25^, we generated 7 chimeras between mPiezo1 and AtPIEZO in which the C-terminus was derived from AtPIEZO and the N-terminus from mPiezo1 (Extended Data Fig. 2). Using an extracellular Myc tag on the N-terminal mouse-derived sequence, we observed that one of the chimeras mPiezo1/AtPIEZO (CH) with 49% mouse and 51% AtPIEZO trafficked to the membrane of HEK P1KO cells (Fig. 3a and Extended Data Fig. 6). The structural elements required for MA current including the pore, anchor and beam is derived from AtPIEZO. Stretch-activated currents (SAC) were observed in 40% of the cell-attached patches recorded from HEK P1KO cells expressing the CH; 76% of patches from mPiezo1-expressing cells revealed SAC. The maximum current elicited (I_max_; Fig. 3c), negative pressure thresholds (Fig. 3f) and P50 values (Fig. 3g) were similar for mPiezo1 and CH (Extended Data Fig. 7). Interestingly, inactivation of SAC from CH-expressing cells was abrogated and currents were maintained throughout the entire 250ms stretch stimulus (Fig.3e vs d; Extended Data Fig. 7). The lower proportion of SAC-expressing patches in CH is consistent with that observed for mPiezo2^22,25^). The reversal potential of SACs mediated by mPiezo1 and CH were similar (Fig. 3h, i; stretch-induced current is shown in brown), consistent with CH being a non-selective cation channel. Thus, the chimera containing the pore-containing C-terminus of AtPIEZO is activated by a mechanical stimulus, suggesting that the native AtPIEZO is indeed a non-selective ion channel in plant cells.

**Figure 3.**
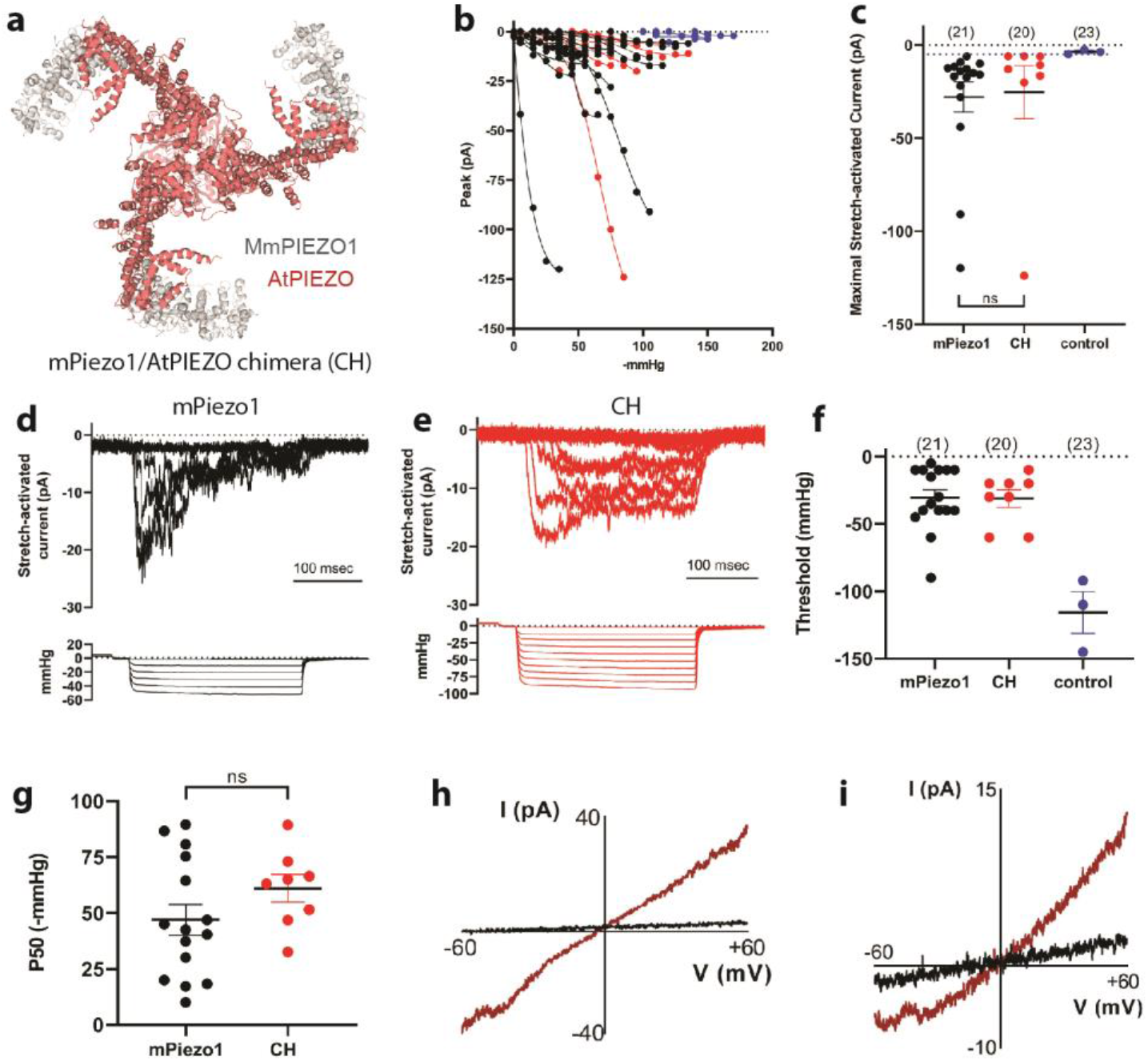
A chimeric channel that includes putative AtPIEZO pore sequences is activated by negative pressure applied to cell-attached patches. **a**, The region of AtPIEZO in the chimera (CH) is highlighted in the mPIEZO1 structure, grey is mPiezo1 sequence (577-1185aa) and red is *AtPIEZO* sequence (1228-2485aa). The first 577 amino acid of mPIEZO1 are not resolved in the structure. **b**, Stimulus-response curves are shown for all mPiezo1 (black) and mPiezo1/AtPIEZO chimera (CH, red) stretch-activated currents (SAC, V_pipette_ = +80mV in cell-attached configuration). Small high threshold responses are occasionally observed in HEK P1KO cells transiently transfected with the empty IRES-GFP vector control (blue). The maximal current observed from vector-transfected cells was −5.4 pA (dotted line) and this value is used as a cutoff for identifying mPiezo1- and CH-mediated SAC. **c**, I_max_ is shown for mPiezo1 (black), CH (red), and control cells (blue). **d** and **e**, SACs recorded in a patch from a cells expressing either mPiezo1 (**d**) or CH (**e**); current amplitudes increase with increasing negative pressure (shown below each family of currents). **f**, The negative pressure (mmHg) at which the first response to stretch is observed (threshold) when patches are challenged with −5mmHg increments (threshold) is plotted. **g**, The pressure producing half-maximal currents (P50, determined using GraphPad Prism) is shown**. h** and **i**, A stretch stimulus eliciting a submaximal response is applied during a voltage ramp protocol in order to record SAC currents between ± 60mV and determine the apparent reversal potential (V_rev_).

We next investigated whether the observed root growth phenotype could be attributed to compromised PIEZO channel activity in root tips challenged with mechanical forces. To accomplish this, we monitored calcium influx in response to mechanical stimulation *in vivo* using a GFP based Ca^2+^ indicator (GCaMP3) expressing transgenic line^26^. First, we applied a localized stimulation by a blunt glass pipette to the root cap of WT plant in increments of 20 μm (Fig. 4a, b). The transient and localized Ca^2+^ signals appeared in the columella cells and LRC cells starting at an indentation of ~60 μm, while peak Ca^2+^ signals were observed at 80 μm of indentation (n=12) (Fig. 4c, d; Supplementary Video 2). At 100 μm indentation and beyond, Ca^2+^ signals propagated between neighboring cells bidirectionally in a manner similar to a wound-mediated response and were not studied here^26,27^ (Supplementary Video 3). We next generated a model in which PIEZO was knocked down specifically in columella cells by using the artificial PIEZO-targeting microRNA driven by the PIN3 promoter^28^ (Fig. 4a). This approach provides an internal control within each root tip: mechanical stimulation-dependent Ca^2+^ responses in columella cells (where AtPIEZO is knocked down) can be compared with neighboring WT LRC cells. Using this strategy, we observed normal responses in LRC cells but significantly reduced responses in the columella cells at 80 μm of indentation (n=9, Fig. 4e-h; Supplementary Video 4). indeed, the area under the curve and the peak of GCaMP3 signals were significantly decreased in the columella but not LRC cells of *piezo* knock-down plants compared to WT plants (4i,j). These data indicate that mechanical stimuli can induce calcium transients in columella cells through AtPIEZO.

**Figure 4.**
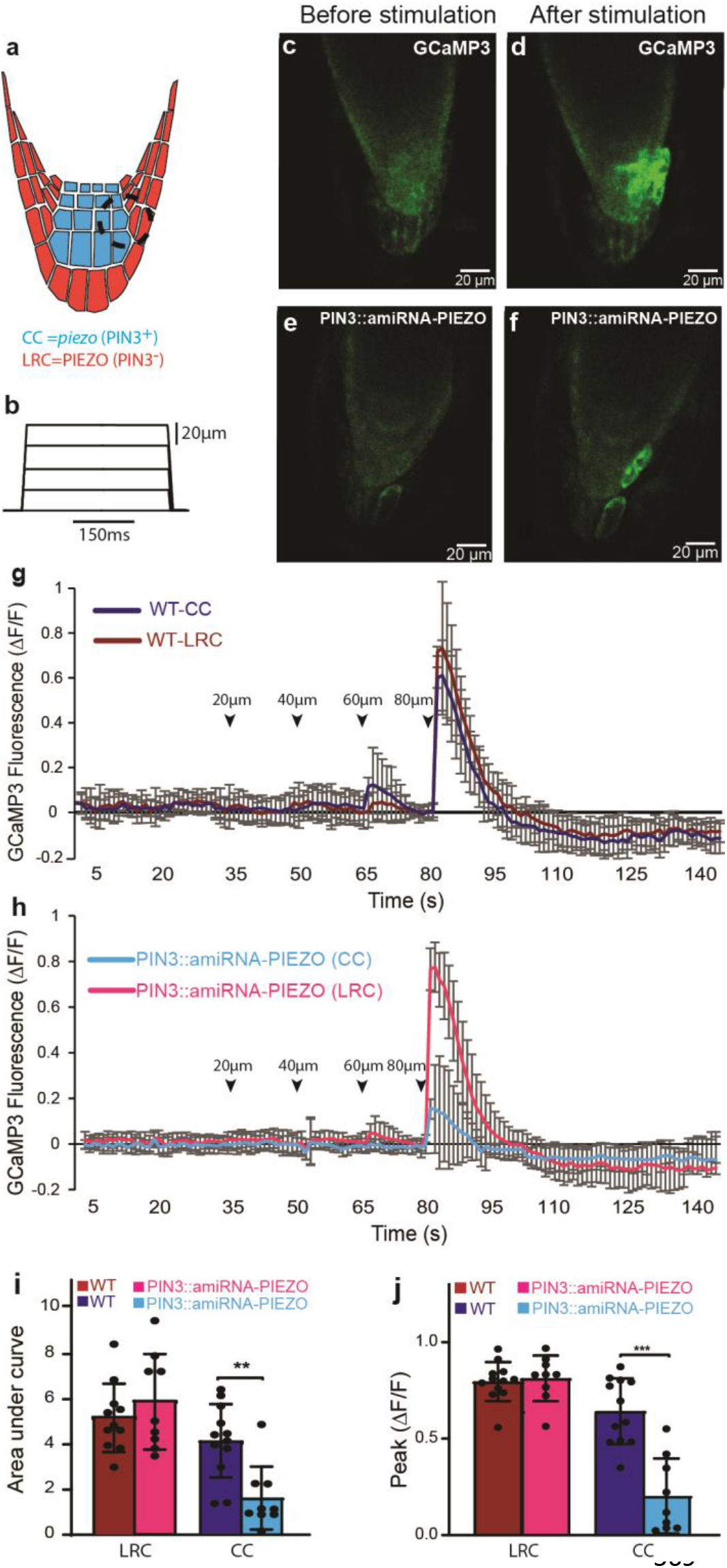
AtPIEZO and calcium influx in response to mechanical indentation to the *Arabidopsis* root tip. **a**, AtPIEZO expression was knocked down in the root cap using an artificial microRNA under the *PIN3* promoter (PIN3::amiRNA-PIEZO) such that *AtPIEZO* is specifically knocked down in columella cells (CC) (blue) but AtPIEZO is still expressed in lateral root cap (LRC) cells (red). Black dashed circle indicates the area of stimulation by the blunt-tip pipette and the region of interest (ROI) for Ca^2+^ signal intensity measurement. **b**, Displacements of the root cap by the stimulating pipette. Mechanical stimulation were initiated 30-35 s after GCaMP3 signal recording; A ramp (1μm/ms) and hold (150 ms) stimulus was applied in 20 μm increments every 15 sec. **c** and **d**, Representative image of the GCaMP3 Ca^2+^ response in a WT root tip before (**c**) and after (**d**) a 80 μm mechanical stimulation (images are from the Supplementary Video 2). **e** and **f**, Representative image of the Ca^2+^ responses in *AtPIEZO* knockdown PIN3::amiRNA-PIEZO before and after a 80 μm mechanical stimulation (images are from the Supplementary Video 4). **g**, Ca^2+^ responses in the LRC cells and columella cells in response to mechanical stimulation in 7 day old seedlings; N=12 (mean±s.d.). Arrowheads indicate the mechanical stimulation (μm). **h**, Ca^2+^ responses after mechanical stimulation in AtPIEZO knockdown PIN3::amiRNA-PIEZO root cap in 7 old days seedlings. Ca^2+^ transients are significantly reduced in columella cells compared to LRC cells; N=9 (mean±s.d.). **i**, Area under curve from starting the 80μm mechanical stimulation until the GCaMP3 signals returned to base line (total of 15s). **j**, Maximum peak of fluorescence in WT and PIN3::amiRNA-PIEZO (*piezo* knockdown) (N=9-12, mean ±s.d.). \*\**P* < 0.01, \*\*\**P* < 0.001.

Plant roots sense physical properties of the soil to either avoid it or penetrate it^3^. Here, we report that *Arabidopsis* PIEZO activity is required for proper root penetration in compacted environments imposing mechanical stresses. PIEZO proteins from numerous animal species are established physiologically relevant MA cation channels^15–17^. We present evidence to suggest that AtPIEZO is functionally conserved as a mechanosensitive ion channel in plant roots. Using calcium imaging we identify at least one cell type in the root cap (columella cells) that requires AtPIEZO to respond to a mechanical stimulus with increased calcium transients. Mutants in other components of calcium signaling pathways such as clm24 (tch2) show similar growth defects to those reported here for *piezo*^29^. The receptor-like kinase FERONIA maintains cell wall integrity through a direct interaction between its extracellular domain and components of the cell wall; it has been proposed to activate a calcium permeable channel whose identity is unknown^30^. AtPIEZO protein might directly alleviate mechanical pressure in columella cells by protecting cell wall integrity and/or by transducing Ca^2+^ signals to other parts of the root such as the elongation zone. Our findings will enable future research to understand the molecular and cellular pathways involved in mechanotransduction within roots. Our results also suggest that other MA ion channels contribute to barrier penetration since the root growth deficits observed in *piezo* mutants are incomplete. In summary, we provide evidence that AtPIEZO acts as a mechanosensitive ion channel in root tips: it has appropriate expression in the root, is required for responses to acute mechanical stimulations and for proper root growth, and it forms an ion channel. These results demonstrate that AtPIEZO mediates a mechanosensory function in *Arabidopsis*, highlighting a conserved function of PIEZOs from plants to mammals.

## Method

### Plant material and growth conditions

*Arabidopsis thaliana* (Col-0) were soil-grown on a 14h light/10h dark cycle (100 μE s^−1^ m^−2^), 70% humidity at 22 °C. For seedling propagation, seeds were surface-sterilized with 70% ethanol for 15 min and washed 5 times with sterile distilled water. Seeds were grown on plates containing half-strength Murashige and Skoog solid medium (0.5X MS) (Sigma-Aldrich, M5524), 0.5 g/L MES hydrate (Sigma-M2933), pH adjusted to 5.7, and then supplemented with 8.5 g/l agar (Sigma-Aldrich, A1296). Plates were placed in a growth chamber vertically or as indicated.

### Hard agar media and barrier plates

First, we poured 160 ml of higher agar concentration in 0.5X MS media into 120mmx120mmx15mm plates. After the plates had cured, we cut the solidified media with a sharp sterile blade into four parts. The first and third segment are removed and replaced with standard MS media containing agar at 8.5 g/L concentration with 5 mm thickness.

### Generation of *AtPIEZO_pro_:GUSPlus* plants

In order to determine tissue specific AtPIEZO expression, we used GUSPlus protein under the *AtPIEZO* promoter. Since, there is an annotated gene close to *AtPIEZO* in the opposite orientation, two different GUSPlus lines containing promoter sizes of 823 bp and 2000 bp were generated. In order to faster screen the transgenic seeds, we first inserted the FAST (fluorescence-accumulating seed technology) cassette^31^ into pCAMBIA-0380 plasmid (CAMBIA), referred to as pCAMBIA-FAST. The FAST cassette was amplified from *VSP2_Pro_:GUSPlus* plasmid^32^ using forward atgttgggcccggcgcgccgagatctTCTAGTAACATAGATGACACC and reverse tggctgcaggtcgacgTCTAGAGGTACCCGGGATCCAGTGTATGTAGGTATAGTAACATG primers. pCAMBIA-380 was cut by *Eco*RI and *BamHI* restriction enzymes (ThermoFisher). Both insert and plasmid were ligated using the Gibson assembly kit (New England Biolabs). The *AtPIEZO* promoters were amplified using forward, ttgttcatgttactatacctacatacactgtggaaagaaagtaaaggattag and reverse, agtagccatgTGGAAACTTTTGTCTTAACG primers and GUSPLUS amplified from *VSP2_Pro_:GUSPlus* plamid using forward, aaagtttccaCATGGCTACTACTAAGCATTTG and reverse gtcagatctaccatggtggactcctcttaaCAATTCACACGTGATGGTG primers. pCAMBIA-FAST was cut using BamHI and HindIII. Both inserts and plasmid were ligated by the Gibson assembly kit. *Agrobacterium tumefaciens* (GV3101) competent cells were transformed with this plasmid. Seeds that expressed red fluorescence protein (RFP) were selected by fluorescence microscopy. The T_3_ generation was used for GUS staining experiments.

### GUS staining and cross sectioning

Transgenic *Arabidopsis* plants expressing β-glucuronidase (GUS) were stained following the protocol described by Jefferson^33^. 5-10 day old seedlings or flowers were collected. Samples were stained with 1 mM X-Gluc (Thermo Fisher Scientific, R0851) in a pH 7.0 phosphate buffer containing 10 mM EDTA, 0.1 mM potassium ferricyanide, 10% (v/v) Triton X-100 at 37 °C overnight. The tissue was destained with serial dilution of 25%, 50%, 75% and 95% (v/v) ethanol. For root cross sectioning, the GUS stained roots were submerged in a box containing Tissue-Tek O.C.T compound (Sakura FineTek USA) and then frozen in liquid nitrogen. Different layers of the root were sectioned at 15 μm thickness and placed on coverslips.

### Generating *piezo* mutants and genotyping

Since homozygote seeds from Salk T-DNA lines were unavailable, we used CRISPR/Cas9 gene editing technology to generate *piezo-FL* (full length deletion) and *piezo-CT* (C-terminal deletion) knockout mutants^34–36^. Both *piezo* mutants were generated in the pNano65 Ca^2+^ indicator background^37^ and these lines and pNano65 WT were used for all experiments except Ca^2+^ imaging experiments. Guide RNA 4 and 6 were used for generating a whole deletion in the *PIEZO* gene, and guide RNA 4 and 5 were used for generating a C-terminal deletion from the LGYL motif to the end of the *PIEZO* gene (1253aa to 2485aa). The target sequences for the whole deletion in *PEIZO* were target 4 CCCTCTGCTCTAGCCGCGTACA and target 6 AGGTGCAATATAGTGAACGAGG (PAM sites were underlined). Targets for the partial mutants were target 4 and target 5 CCCAGGTGTAGAATGTCATACT. Primers for genotyping: Piezo-GT1 TCTGCCACATTCCCACTCAG, Piezo-GT2 GGTTTAGCCATTTCTCGGCG, Piezo-GT3 CGGGAGTGTTGGCTTGGTAT, Piezo-GT4 CCGCCACGTAAGTTAGCTCT. The *piezo-FL* mutants were genotyped with the primer pair Piezo-GT1 and Piezo-GT2, which generates a PCR product of 1300 bp in mutants. This primer pair could not amplify WT genomic DNA due to the large size of the fragment. To determine zygosity of *piezo-FL* mutants, we used the Piezo-GT1+ Piezo-GT4, which amplifies a 949bp product from WT DNA, but could not amplify a band in the homozygous mutant. For the C-terminal deletion mutant *piezo-CT*, we used Piezo-GT1 and Piezo-GT3, which would generate a fragment of about 1000bp in mutants. PCR of Piezo-GT1+ Piezo-GT4 was performed for determine the zygosity of *piezo-CT* as well.

### Generating piezo knockdown mutant

We used atMIR390 microRNA^38^ to knockdown PIEZO in columella cells. We synthesized atMIR390 microRNA that included the *PIEZO* targeting sequence of TGCAGTTGCTCGGTCTTCCGA and amplified using the forward TTTTTGTCCCTTCAAGTATAGGGGGGAAAAAAAGGTAG and reverse tcttaaagcttggctgcaggGAGACTAAAGATGAGATCTAATC primers. *PIN3* promoter was amplified using the forward cggcgcgccgaattcccgggAATTTTATTGCATATAGTGTGTTTATTAAATG and reverse TTTTTTCCCCCCTATACTTGAAGGGACAAAAATGGAAAAC primers. pCAMBIA-1380 plasmid was cut by *BamHI* and *SalI* restriction enzymes, and the cut plasmids and both fragments were ligated using the Gibson assembly kit. *Agrobacterium tumefaciens*, GV3101 competent cells were then transformed by this plasmid. GCaMP3 plants were transformed using *Agrobacterium tumefaciens* bacteria. MS media containing 10mg/l Hygromycin B was used for screening the transgenic seeds. Two different independent T3 transgenic lines used for Ca^2+^ imaging.

### Quantitative RT-PCR

The upper roots of the *Arabidopsis* plant grown either inside or on top of the surface of the MS media were harvested and total RNA was extracted using the RNeasy plant mini extraction kit (Qiagen). 1 μg of the total RNA was copied into cDNA with The SuperScript III First-Strand Synthesis according to the manufacturer’s instructions (ThermoFisher). Quantitative RT-PCR was performed on 50 ng of cDNA in final volume of 10μl according to the SsoAdvanced Universal SYBR^®^ Green supermix (BioRad). Ubiquitin-conjugating enzyme (*UBC21*) At5g25760 was used as a reference gene. Four biological replicates, which were mixture of 3-5 plants, were used for each experiment. Primers used were: *UBC21* (At5g25760) forward CAGTCTGTGTGTAGAGCTATCATAGCAT, reverse AGAAGATTCCCTGAGTCGCAGTT, *AtPIEZO* (AT2G48060) forward ACGCTCTGATATCCAAATGGT, and reverse ACTTCATCCGTCTGATCCTC.

### Cloning *AtPIEZO*

The full length of 7455 bp of *AtPIEZO* were amplified in two parts from *Arabidopsis* total RNA extracted from roots and leaves. The *AtPIEZO* gene was cloned into *pCDNA3-1 IRES2-eGFP Zeo+ (AtPIEZO^GFF^*), a mammalian expression vector. We also generated a codon-optimized *AtPIEZO* for expression in mammalian cell culture (HEK P1KO cells).

### Generating Myc tagged plasmids

The site of Myc tag insertion were chosen based on homology alignments between mPIEZO1 and AtPIEZO and the tag shown to work for these experiments in the extracellular loop of mPIEZO1. The position of the Myc tag is highlighted in Extended Data Fig. 2. The sequence of Myc tag (GAACAAAAACTTATTTCTGAAGAAGATCTG) were insterted in the primers. The Myc tag insertion was cloned into *pCDNA3-1 IRES2-eGFP Zeo+* (*AtPIEZO ^GFP^*). Myc tag 1-3 were inserted in codon optimized *AtPIEZO* and Myc 4 and 5 inserted in the native codon of *AtPIEZO*. All plasmids were generated by the Gibson assembly kit (NEB) according to the manufacturer’s instructions. The sequence of the primers in the following table:

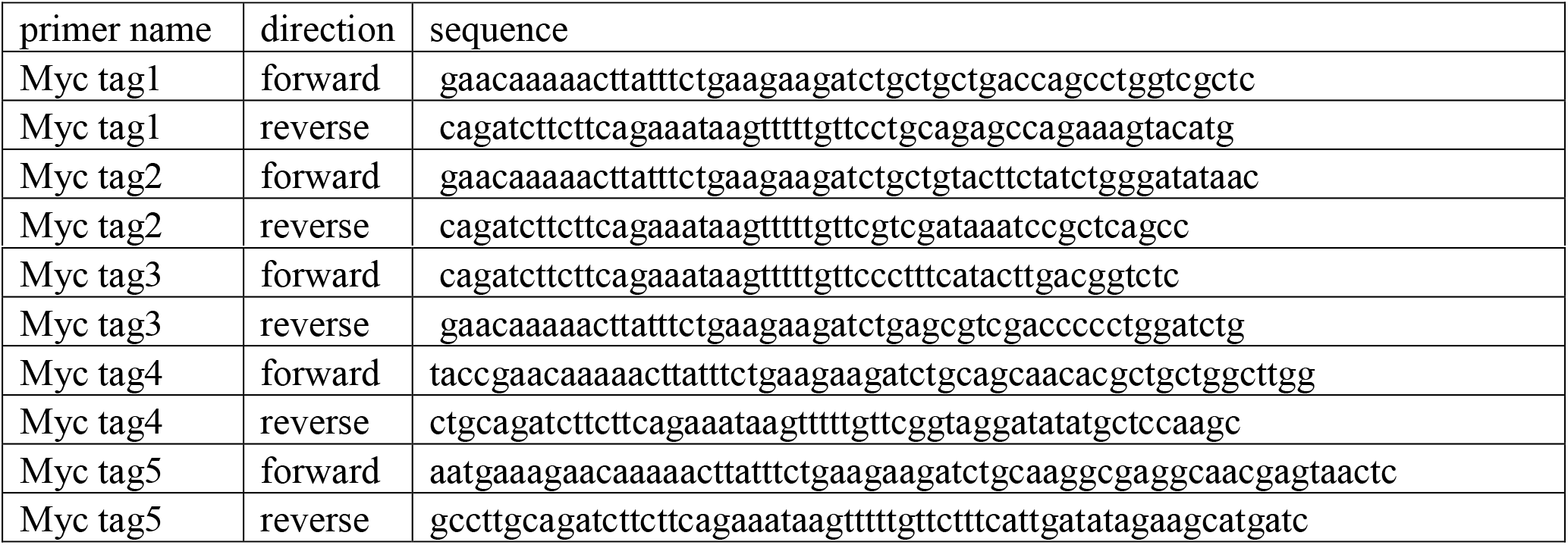

### Generating Chimeras

The chimeras between mPiezo1 and AtPIEZO were generated by swapping specific regions highlighted in Extended Data Fig. 2. The N-terminus of all chimeras was derived from mPiezo1 and C-terminus was derived from a mammalian codon optimized AtPIEZO. The junction between chimeras was chosen based on conserved amino acid regions between mPiezo1 and AtPIEZO. We amplified the fragments using Q5 polymerase (NEB). *pCDNA3-1 IRES2-eGFP Zeo+* plasmid was cut by *BamHI* and *EcoRI*, followed by ligation of plasmid and fragments using the Gibson assembly kit (NEB). The ligated plasmids were transformed into XL-gold competent cells and colonies screened. All positive chimera clones were verified by full-length DNA sequencing.

### Electrophysiological characterization of Arabidopsis PIEZO (AtPIEZO), mPiezo1 and the chimera

HEK P1KO cells were transiently transfected with AtPIEZO, mPiezo1, mPiezo1/AtPIEZO chimera (CH), or vector using Lipofectamine 2000 per manufacturer’s instructions and allowed to settle on 12 mm diameter PDL-coated coverslips as previously described^21^.

Electrophysiological recordings were made 2-3 days after transfection using cell-attached patch clamp methods to determine their responsiveness to membrane stretch using a High Speed Pressure Clamp HSPC-1 (ALA Instruments)^22,39^. Pipettes had resistances of 1.4 – 2.6 MΩ when filled with 130 mM NaCl, 5 KCl, 1 CaCl_2_, 1 MgCl_2_, 10 TEA-Cl, and 10 HEPES (pH 7.3 with NaOH). Cells were exposed to in a high KCl bath solution during recordings to depolarize the membrane potential (140 KCl, 1 MgCl_2_, 10 glucose, 10 HEPES (pH 7.3 with KOH)). Negative pressure steps (250 msec in duration) were applied after 6 sec at +5 mmHg followed by 20ms at 0 mmHg; steps of increasing negative pressure in −5 mmHg increments were applied every 15 sec. For SAC recordings, the Multiclamp700A feedback resistor used was either 5 or 50 GΩ depending on the size of the currents. To determine the apparent reversal potential (V_rev_), a stretch stimulus eliciting a submaximal response was applied during a voltage ramp protocol in order to record SAC currents between ± 60mV. The apparent V_rev_ determined from this voltage ramp protocol was validated in the same patch with a typical voltage step protocol. Whole cell recordings were made as described^16^ for AtPIEZO-transfected cells 2-3 days after transfection.

### *In vivo* Ca^2+^ imaging

*A. thaliana* plants expressing genetically-encoded Ca2+ (GCaMP3) that obtained from Dr. Edward E. Farmer (university of Lausanne) were used for *in vivo* imaging. Images were collected with a Nikon Instruments A1R+ confocal mounted onto an inverted Ti-E microscope. An S Plan Fluor ELWD 20x objective NA 0.45 was used to acquire images at 1 frame/sec 1024 x 512 scan area (frame rate 0.946 msec/frame), 0.62 microns/pixel, pinhole 1.4AU, laser power 0.2 microwatts out of the objective. A Coherent 488nm solid-state laser was used for excitation, with a Chroma 525/50 emission filter. Nikon Elements software was used for timelapse intensity measurements. GCaMP3 imaging was recorded 30-35 s before applying mechanical stimulation. A day before Ca^2+^ imaging, 5-7 day old seedling was transferred onto a 60mmx24mm coverglass covered by 1-2mm of MS media in 0.6% low melting agarose (IBI scientific). The GCaMP3 plants were excited using a mercury lamp, 488nm laser and emission filter of 525/50nm with andore 897EMCCD camera. The GFP signals of several regions of interest (ROI) such as columella cells and lateral root cap cells analyzed using the NIS-Elements imaging software. Representative images are shown in Figure 4, after adjusting brightness and contrast for clarity in publication. We used (ΔF/F), the equation ΔF/F = (F - F0)/F0 for analyses the fluorescence changes. F0 is baseline fluorescence that calculated from 10s before stimulation and F is fluorescence of the recording. We estimated the area under curve using the equation (y1+y2)/(2*(t2-t1)), where y is the value of (ΔF/F) and t is the each time point of GCaMP3 recording.

### Mechanical stimulation to root

For plant *in vivo* Ca^2+^ measurements, mechanical stimulation was achieved using a fire-polished glass pipette (tip diameter 10-12 μm) positioned at an angle of 80° to the recorded cells. Downward movement of the probe was driven by a Clampex controlled piezo-electric crystal microstage (E625 LVPZT Controller/Amplifier; PhysikInstrumente). The probe had a velocity of 1 μm.ms-1 during the ramp segment of the command for forward motion and the stimulus was held for 150 ms before releasing the stimulus. To assess the mechanical responses of a cell, the probe was first placed as close to the cell as possible (this distance could vary from plants to plants). We optimized the mechanical stimulation and found that 4 series of mechanical steps in 20 μm increments in every 15s lead to a transient and local Ca^2+^ responses. Longer or more steps led to Ca^2+^ fluxes traveled bidirectionally into both sides of the region being poked resemble of damage/wounding Ca^2+^ signaling.

### Statistics

All results in main figures and extended data with error bars are represented as mean ± s.d. according to standard methods using Microsoft Excel or GraphPad Prism. The *P* values were generated with Student’s one-tail unpaired *t*-tests. For qRT–PCR experiment, four technical replicates were used (three technical replicates). The biological replicates were indicated as ‘*n*’ in the figure legends.

## Acknowledgements

We would like to thank Dr. Simon Gilroy, and Dr. Edward E. Farmer and Dr. Elliott Meyerowitz for seeds. We also thank Dr. Kathryn R. Spencer for helping *in vivo* Ca^2+^ imaging. Dr. Elizabeth S. Haswell, Haley DeGuzman, Tess Whitwam, Shang Ma, Ivan Radin, Kara Marshall, Yunxiao Zhang, Swetha Murthy and Viktor Lukacs for discussion and critical reading of the manuscript.

## Funding acknowledgments

Early and advanced mobility fellowship from Swiss National Science Foundation (S.A.R.M.). YZ was supported by NIH GM114660, and CG was supported by a scholarship from the China Scholarship Council.

## Author contributions

S.A.R.M., A.E.D. and A.P. conceived and designed experiments and wrote the manuscript. A. E. D. and W.Z.Z. recorded and analyzed electrophysiology data. C. G. and Y. Z. generated the CRISPR-Cas9 *piezo* mutants. A. C. performed live labeling. K. D. and D. G. performed cloning and generated transgenic plants.

## Corresponding authors

To whom correspondence may be addressed.

---

**Extended Data Figure 1.**
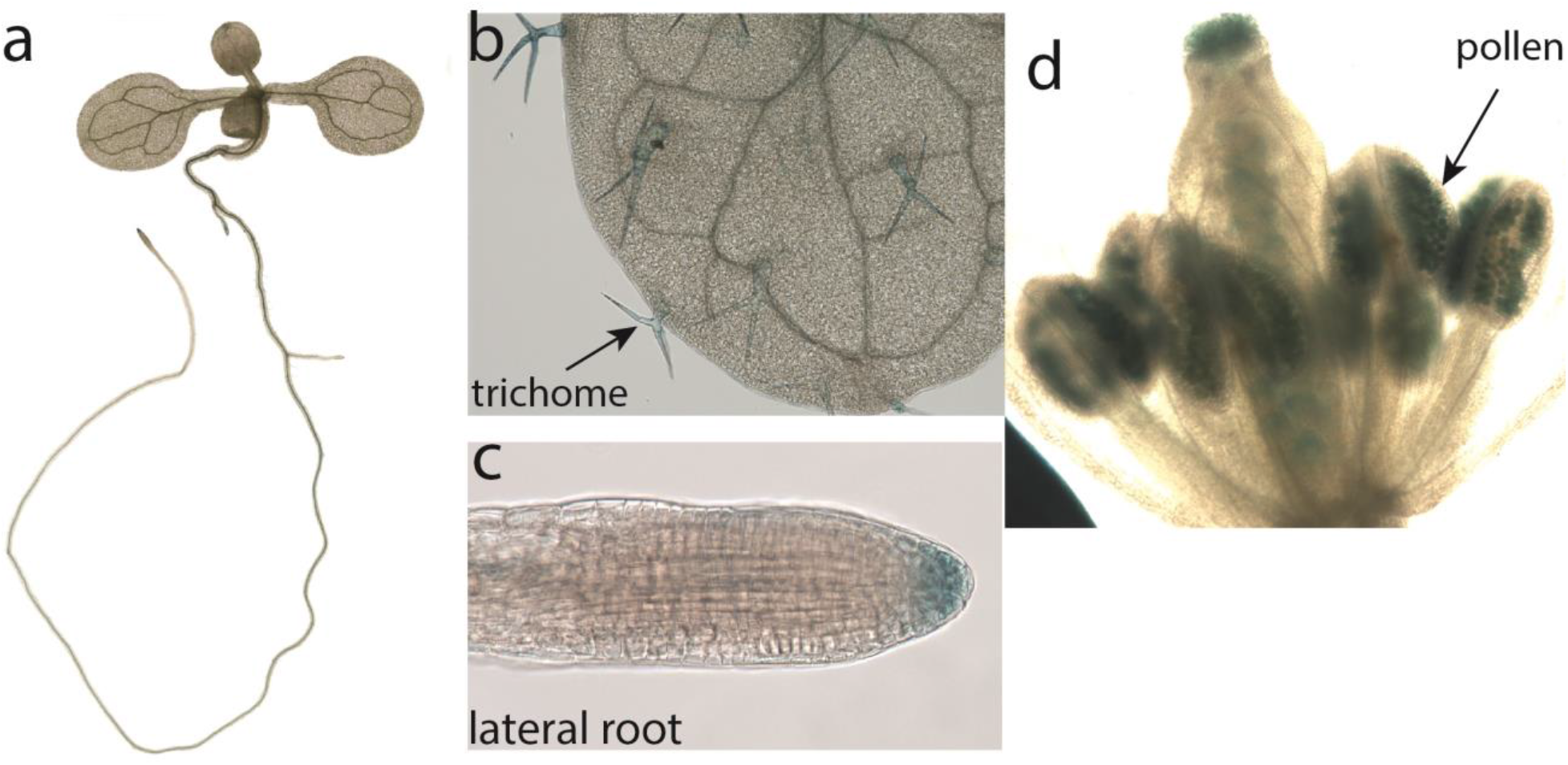
Representative images of the expression pattern of *AtPIEZO* promoter activities in *AtPIEZO::GUSPlus* reporter line. **a**, The expression pattern of *AtPIEZO* promoter activity in the line with the 823 bp *AtPIEZO* promoter (*AtPIEZO _(short)_::GUSPlus’*). **b-e**, The expression pattern of *AtPIEZO* promoter activity in the line with 2000 bp *AtPIEZO* promoter (*AtPIEZO _(long)_::GUSPlus*) in leaf that indicates *AtPIEZO* expression in trichome, laterial root cap, flower and pollen.

**Extended Data Figure 2.**
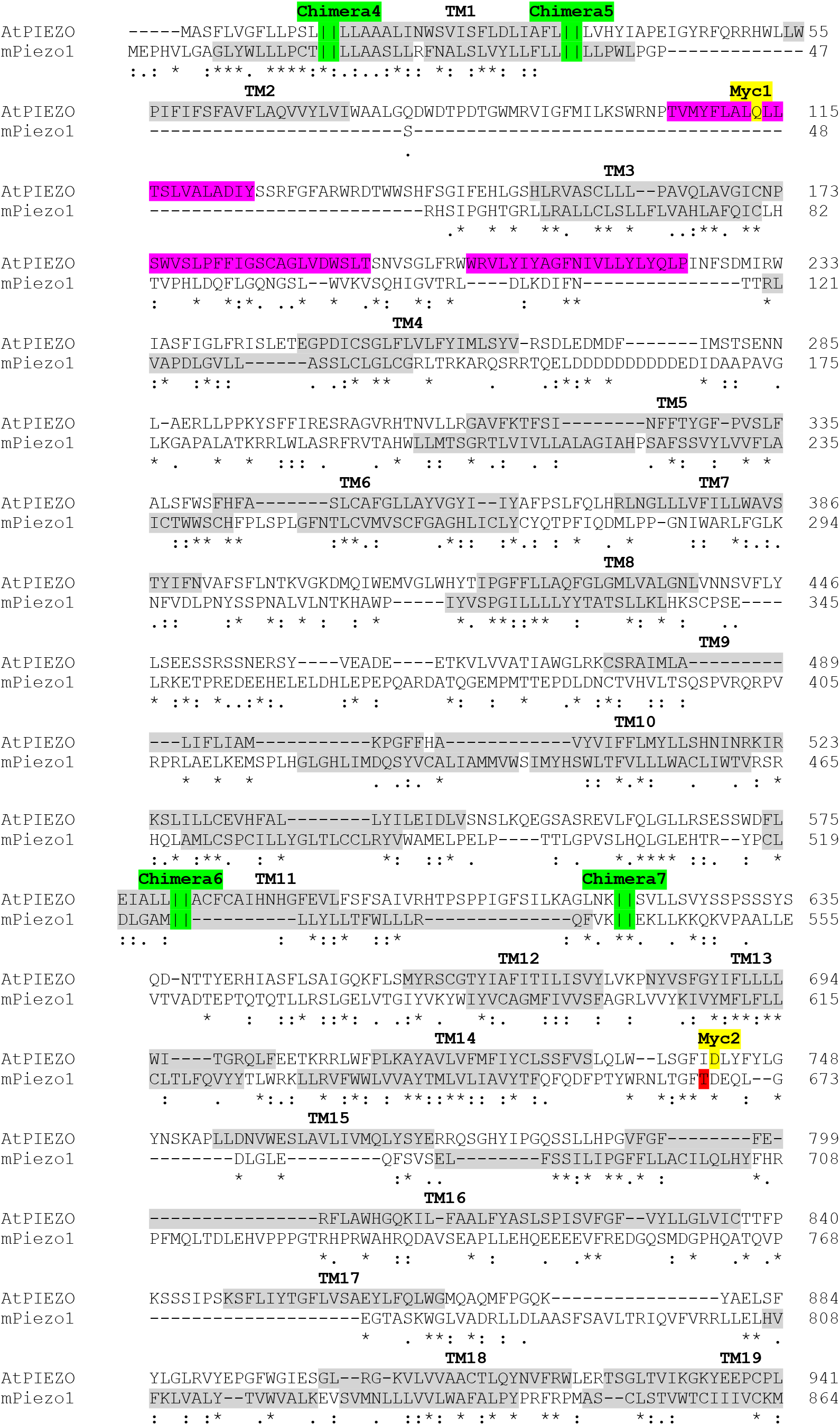
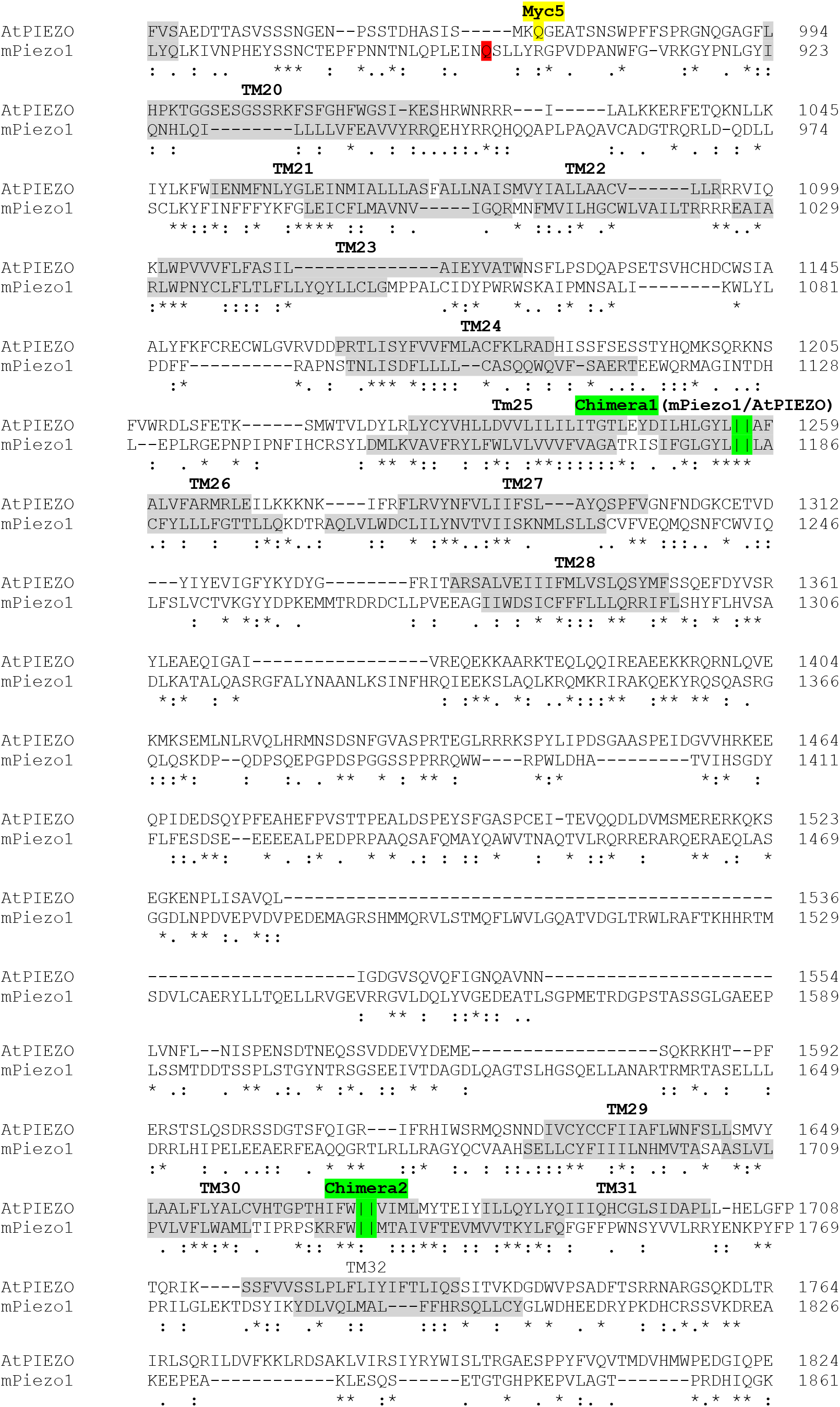
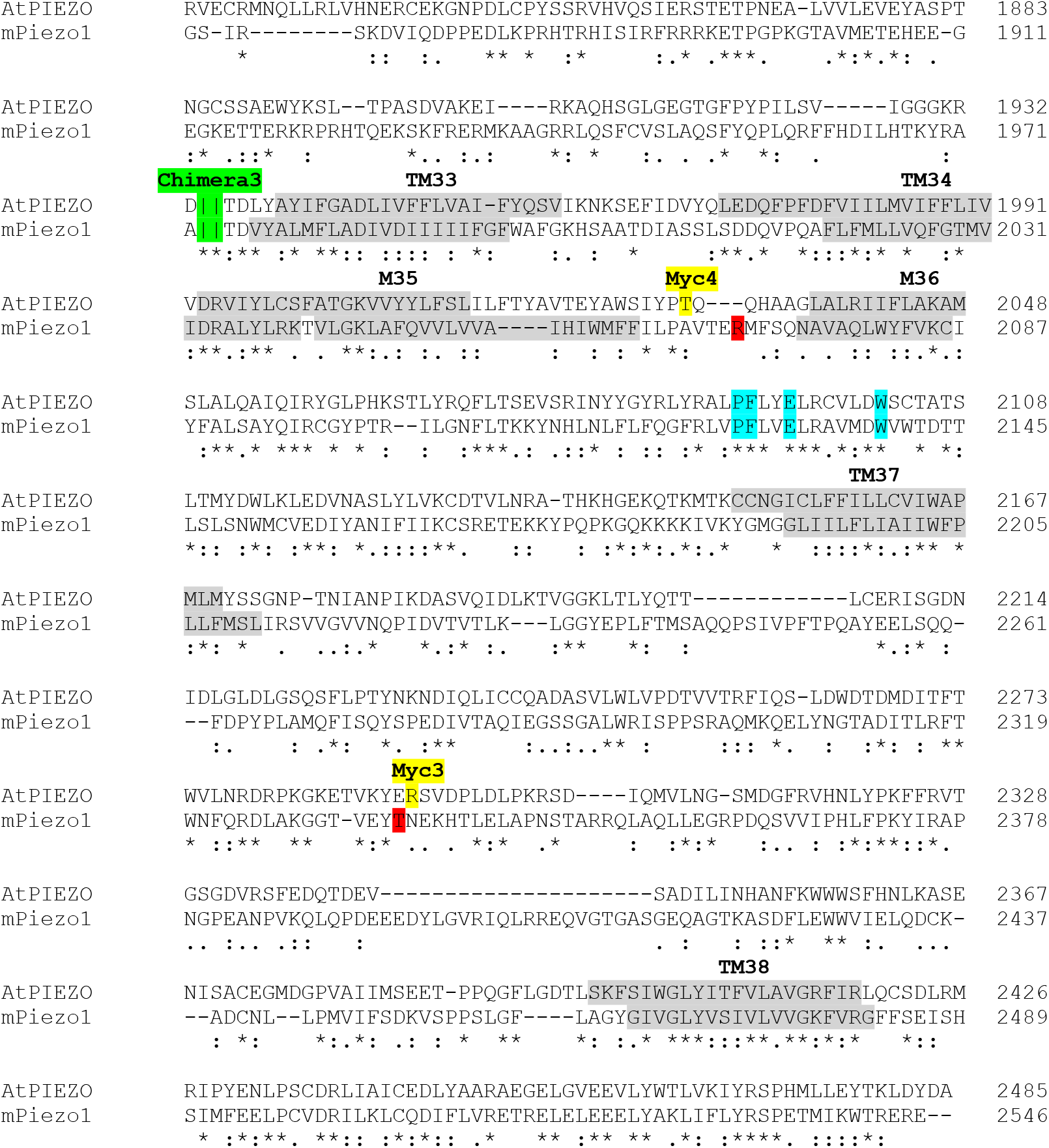
Alignment between *Arabidopsis* PIEZO and mouse Piezo1 highlights the residues of interest. A multiple sequence alignment between mPiezo1 and AtPIEZO was generated using ClustalW2 (http://www.ebi.ac.uk/Tools/msa/clustalw2/). The transmembrane topology prediction for TM1-TM14 was obtained using on TOPCONS software (http://topcons.cbr.su.se/) and the topology from TM15 to TM38 was derived from the structure of mPiezo1^23^. Residues highlighted in grey indicate the transmembrane domain. Residues in pink are transmembrane domains predicted to be only in AtPIEZO, but not mPiezo1. Note that there is higher homology between mPiezo1 and AtPIEZO in the regions where the structure of mPiezo1 is resolved. Residues highlighted in green indicate the junction between mPiezo1/ and AtPIEZO in the chimeras. Residues highlighted in red indicate the position of the Myc tag on mPiezo1^22^. Residues highlighted in yellow, indicate the position of the Myc tag on AtPIEZO. The PFEW motif highlighted in blue conserved among plants, mammals and protozoa^40^.

**Extended Data Figure 3.**
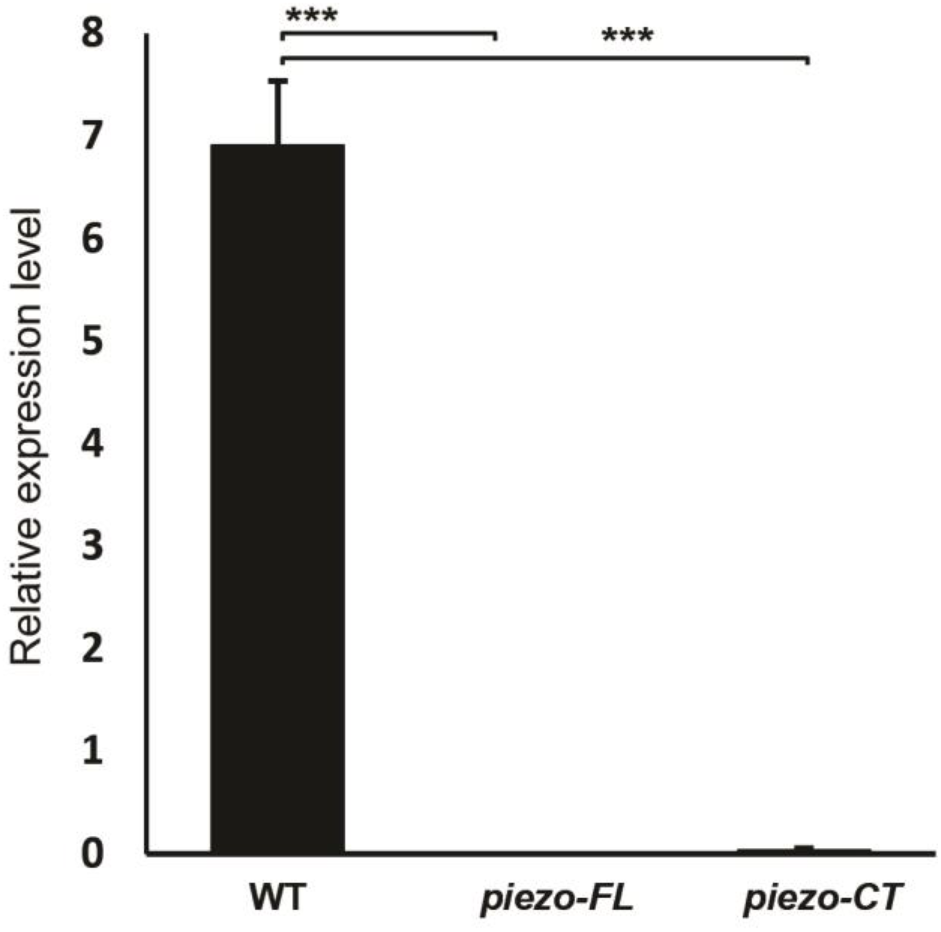
*AtPIEZO* transcript level in roots of WT and mutant plants. qRT-PCR was performed on samples harvested from root and leaf from 4 different plants. \*\*\**P* < 0.001, N=4 (mean ± s.d.).

**Extended Data Figure 4.**
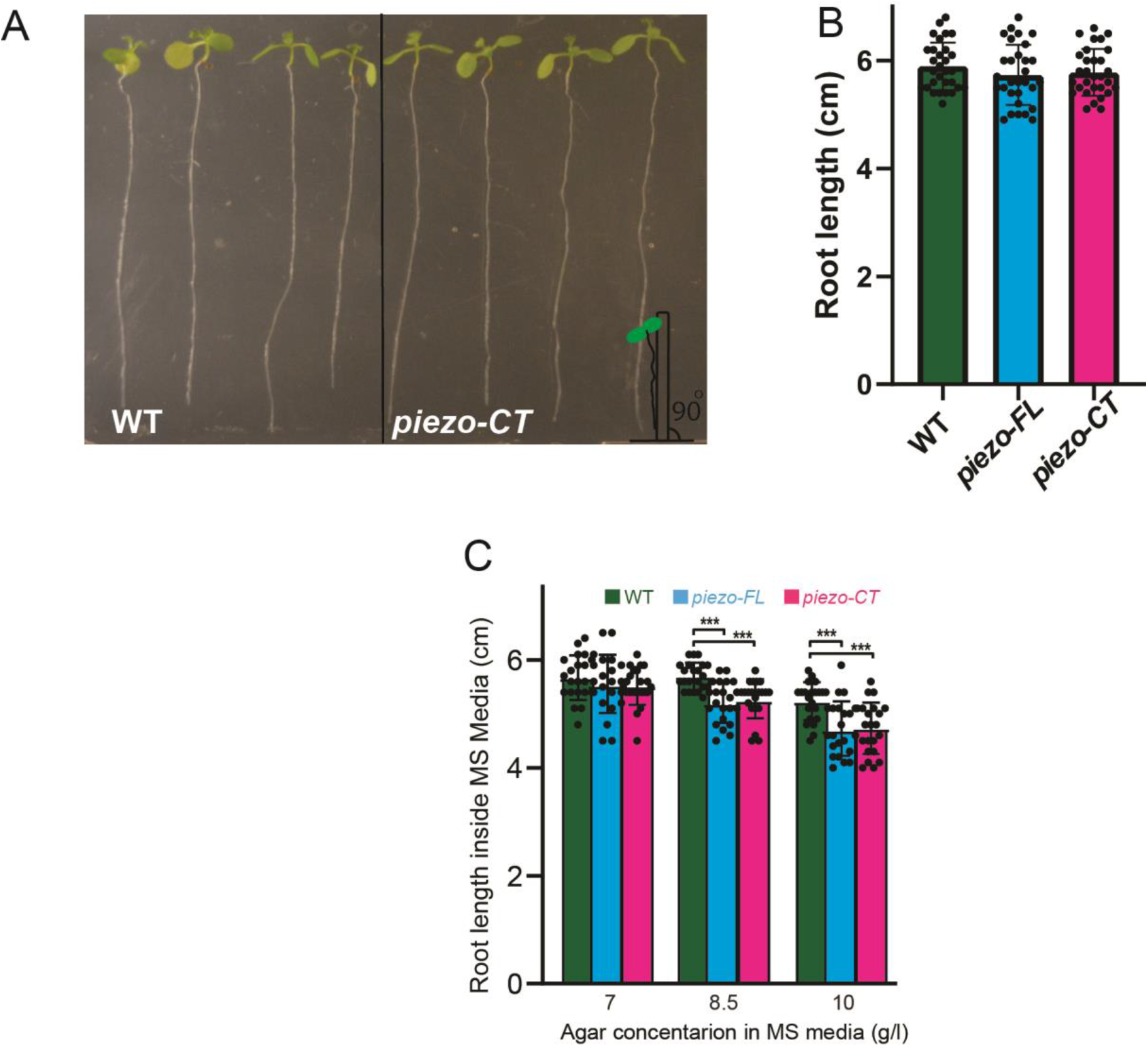
Root length of *piezo* mutants. **a**, Representative image indicating the root lengths of *piezo* mutants when grown on top of MS media in plates tilted at a 90° angle. **b**, Root length of WT and both *piezo* mutants *piezo-FL* and *piezo-CT* (n=30). **c**, Root length of *piezo* mutants when grown inside the MS media containing the indicated agar concentrations in plates positioned at a 60° angle. Data shown are for roots growing inside the MS media. \*\*\**P* < 0.001 (N=23, mean±s.d.).

**Extended Data Figure 5.**
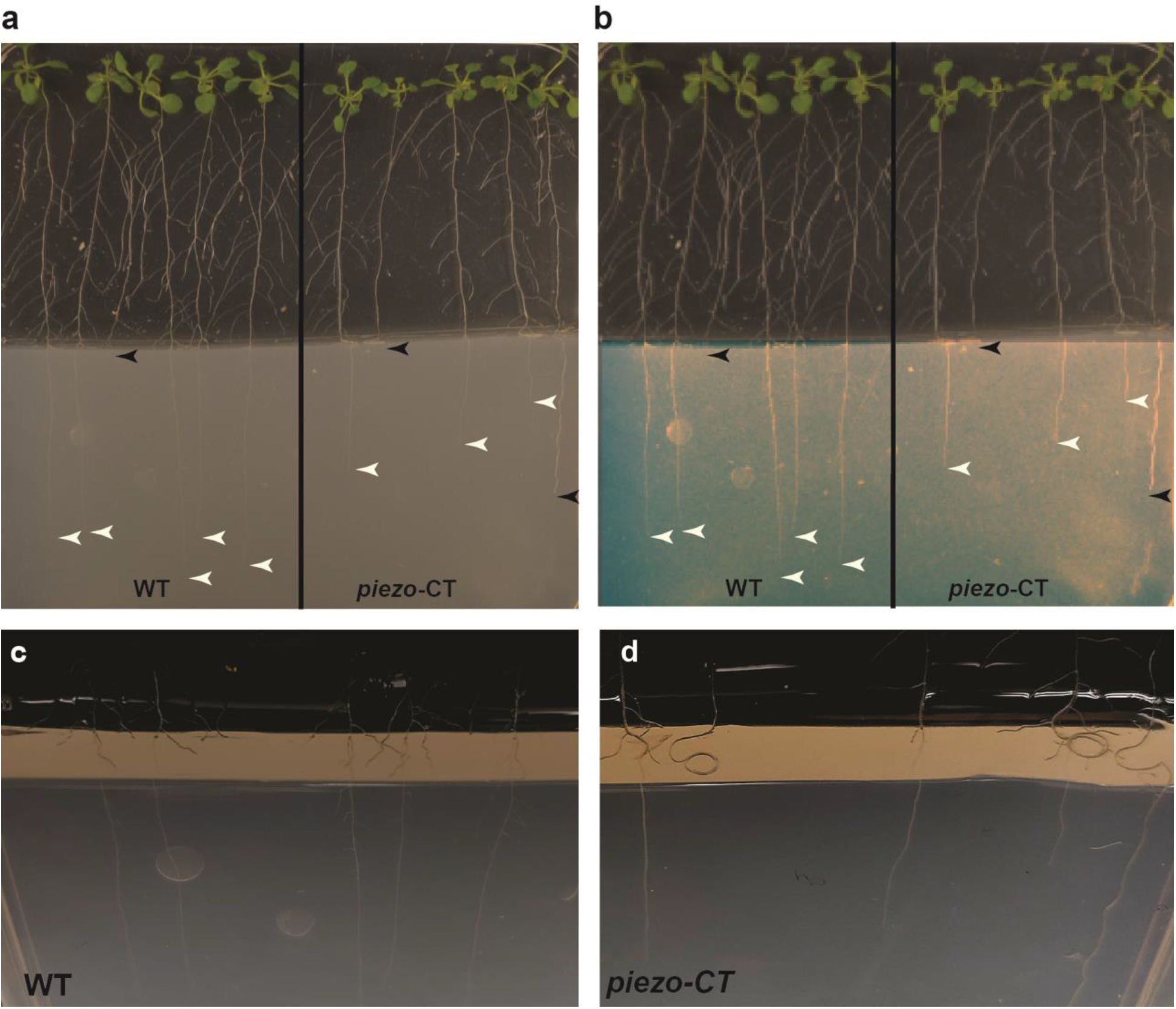
*piezo* mutants are defective at penetrating a hard barrier. **a**, Original image of the plant root that was challenged by barriers from Fig. 2e. **b**, Same image with adjusted light exposure for better visibility of roots grown inside the MS media. Black arrowheads indicate roots within the MS media or at the barrier interface, white arrowheads indicate root growth on the surface of MS media. **c** and **d**, The barrier is imaged at an angle that enables visualization of the root tips at the 5 mm wide barrier (brown; same data shown in panel A). **c**, Usually WT roots are observed to grow fairly straight through the harder barrier. **d**, Some of *piezo-CT* roots are observed to also form swirls at the barrier.

**Extended Data Figure 6.**
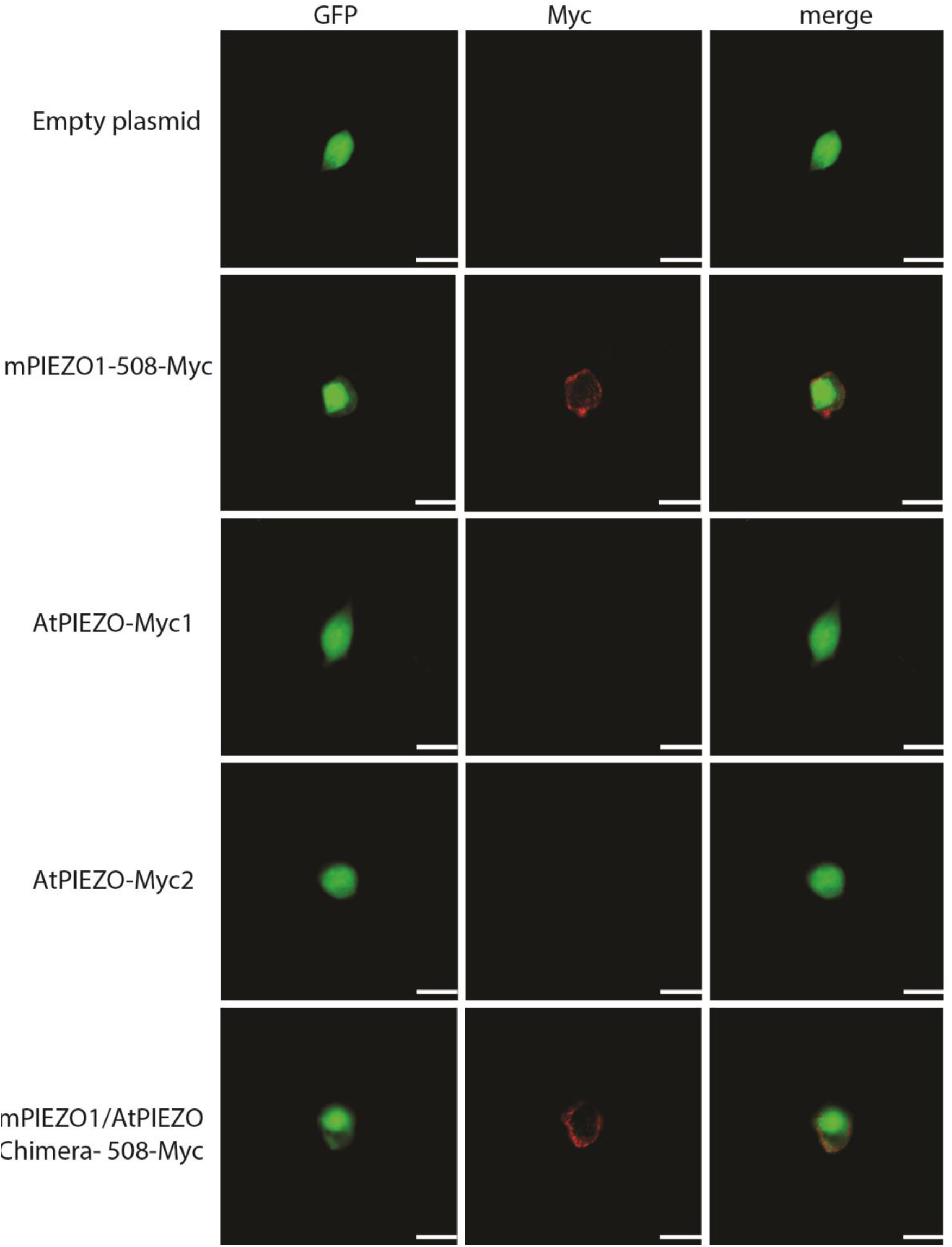
Myc tag staining of AtPIEZO and chimera. Representative images of non-permeabilized staining using an anti-Myc antibody (red) in *AtPIEZO-myc* -ires-GFP transfected cells, mPIEZO1-508-Myc (Myc tag located after 508 amino acid) and mPIEZO1/AtPIEZO chimera containing 508-Myc. mPIEZO1-508-myc used as positive control^22^. Scale bar is 20 μm.

**Extended Data Figure 7.**
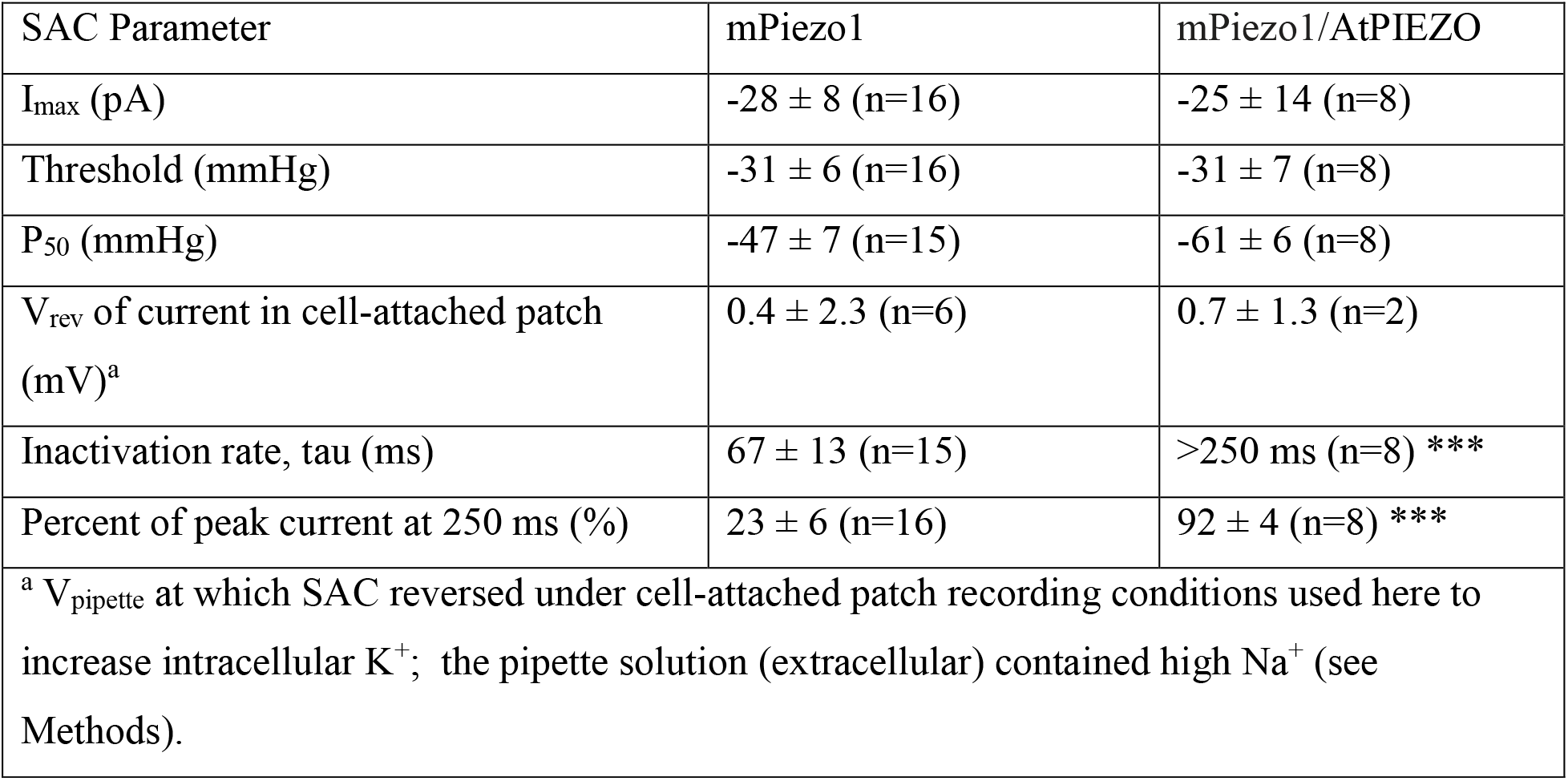
Electrophysiological characterization of mPiezo1 and mPiezo1/ AtPIEZO chimera.

**Supplementary Video 1.** *piezo* mutant poorly penetrate into hard MS media. Seeds of WT and *piezo-CT* mutant plated on the surface of agar 2 cm above the barrier (12 g/l agar in MS media).

**Supplementary Video 2.** Mechanical indentation causes Ca^2+^ responses in the lateral root cap cells and columella cells in the WT expressing GCaMP3. Four mechanical stimuli were applied to the root cap beginning at 30s and followed in increasing increments of 20μm at 15s intervals.

**Supplementary Video 3**. Extensive mechanical stimulation (100 μm) lead to wound/systemic Ca^2+^ fluxes that travel in both directions from the stimulation site. Five mechanical stimuli were applied to the upper root beginning of 25s followed in increasing increments of 20μm at 15s intervals. At the 100 μm of mechanical stimulation, Ca^2+^ responses travel bidirectionally.

**Supplementary Video 4.** Mechanical indentation causes Ca^2+^ responses only in the lateral root cap cells in *piezo* knockdown (PIN3::amiRNA-PIEZO) mutant expressing GCaMP3. Four mechanical stimuli were applied to the root cap beginning at 30s and followed in increasing increments at 15s intervals.

